# Regional ecological variation drives isotopic niche divergence in Pacific nautiloids

**DOI:** 10.64898/2026.04.29.721722

**Authors:** Job Lukas Veloso, Gregory Barord, Frederick Dooley, Peter Ward

## Abstract

Nautiloids, the only surviving externally shelled cephalopods, persist in isolated Indo-Pacific reef slopes despite life-history traits that limit dispersal and recovery. Yet the ecological basis of their persistence remains poorly understood. Here, we compare carbon (δ^13^C) and nitrogen (δ^15^N) isotope values from seven nautiloid populations (including *Nautilus* and *Allonautilus*) spanning the Pacific. Isotopic niches varied strongly among locations, but only weakly among species, suggesting that geographic context rather than phylogenetic identity is the primary driver of trophic differentiation. Populations from the Bismarck Sea and American Samoa exhibited elevated δ^15^N, consistent with regional nutrient cycling and nitrogen fixation, whereas Great Barrier Reef (GBR) nautiloids displayed unusually broad δ^13^C ranges linked to possible reef-derived carbon subsidies. These results reveal how local oceanography and resource availability shape isotopic niches in long-isolated populations, providing a framework for understanding both the ecological resilience and evolutionary divergence of ancient cephalopods in modern oceans.

## Introduction

Cephalopods occupy diverse marine ecosystems and play pivotal roles in food webs, yet their ecological dynamics are shifting under global change[1–4]. Unlike most coleoids, nautiloids are slow-growing, late-maturing, and geographically restricted, traits that heighten their vulnerability to environmental and anthropogenic pressures[5–12]. Extant species of *Nautilus* and *Allonautilus* are confined to deep fore-reef slopes (100–800 m) across the Indo-Pacific, where their populations are geographically isolated by oceanic barriers and depth limits[13–15]. This isolation, coupled with recent species discoveries[16], suggests complex ecological and evolutionary histories, but the trophic roles of nautiloid populations remain poorly resolved[17,18].

Stable isotope analysis (SIA) offers a powerful lens into ecological interactions by integrating diet and habitat use over time[19–22]. Although isotope values of some nautiloid species have been reported[23,24] these studies have been regional and descriptive. We lack a comparative framework spanning multiple species and locations, which is needed to test whether isotopic niches are structured primarily by phylogenetic identity or by geographic context.

Addressing this gap is important because nautiloids represent the only surviving externally shelled cephalopods, a lineage with deep evolutionary roots. Their persistence across fragmented habitats raises broader questions about how environmental context shapes ecological niches, how geographic isolation influences evolutionary divergence, and how ancient lineages adapt to modern oceans.

Here, we test these ideas across seven populations of *Nautilus* and *Allonautilus* spanning the Pacific. By combining δ^13^C and δ^15^N analyses with Bayesian niche metrics, we assess the relative roles of phylogeny and geography in shaping trophic ecology. Our results provide new insights into niche differentiation, ecological resilience, and evolutionary divergence in one of the oldest surviving cephalopod lineages.

## Materials and Methods

### Sample collection

Tissues were collected from different nautiloid populations across the Pacific for stable isotope analyses. In these sites, weighted traps were deployed between 200 and 400 m depths using a similar method to that used in Vandepas 2016. The weighted traps were made of metal interlinks and approximately measured 2 m x 1 m with a double entry. Raw chicken and canned tuna were used as bait. From each of these sites, captured nautiloids were nonlethally sampled for both shell and tissue. Soft tissue samples (∼ 500 mg) were collected by snipping the tips of a single nautiloid tentacle. The soft tissue samples were then placed in small vials with 95% ethanol.

Additionally, apertural shell snips were obtained and placed in dry vials in air. All samples were returned to the University of Washington, Seattle, WA, USA, for processing.

### Bulk stable isotopes analysis of δ13C and δ15N

Tissue samples were placed in an oven to dry at approximately 60°C for 48 hours. Dried tissues were each sectioned into smaller pieces (∼0.4 mg). Samples were weighed using a microbalance and wrapped in tin capsules. The concentration and isotopic composition of carbon and nitrogen in the samples were measured with a Costech^™^ ECS 4010 Elemental Analyzer coupled to a Thermo Finnigan^™^ MAT253 continuous flow isotope ratio mass spectrometer in IsoLab at the University of Washington. Combustion was carried out at 1000°C with a 10 mL pulse of O_2_. The result of the combustion are gases that are passed through a reduced copper column maintained at 700°C. A magnesium perchlorate trap was then used to remove water from the gas stream, after which the gases were separated via gas chromatography and fed into the mass spectrometer via a Thermo Finnigan Conflo III.

Raw isotopic data were corrected using a two-point calibration with three in-house standards: two glutamic acids and dried salmon, which are calibrated against international reference materials NBS19, LSVEC, IAEA-N-1, USGS32, USGS-40 and USGS-41. Nitrogen isotopic data are reported in delta notation relative to air. Carbon isotopic data are reported in delta notation relative to Vienna Peedee Belemnite (VPDB).

### Data analyses

Linear regression was used to assess the relationship between δ^13^C and δ^15^N and their correlation with shell diameter to detect potential shifts through ontogeny. Our analyses involved ANOVA or Kruskal–Wallis H tests, followed by pairwise multiple comparisons using Tukey’s HSD test or Mann–Whitney U test [25]. All tests were conducted with a significance level of α = 0.05. We used recent metrics based on a Bayesian framework (Stable Isotope Bayesian Ellipses in R: SIBER [26] to analyze stable isotopic niche widths in different groups.

Statistical analysis, computations, and graphs were conducted utilizing R (version 4.3.2), PAST 4.15[27], and MS Excel (version 16.81).

### Ethics Statement

This study did not involve endangered or protected species at the time of sampling. All sampling was conducted non-lethally, and individuals were released following tissue collection.

Research in the Philippines was conducted in collaboration with the University of San Carlos; no collection permit was required as animals were not collected. Research in Australia was conducted under permits from the Great Barrier Reef Marine Park Authority and approved by the University of Queensland Animal Ethics Committee. Research in Fiji was conducted under permit from the Department of Fisheries. Research in American Samoa was conducted under permit from the Department of Marine and Wildlife Resources. This study did not involve human participants, human tissue, or human data.

## Results

### Values of δ^13^C

We observed a significant difference in δ^13^C values among nautiloid populations (F_6,60_ = 14.58, p < 0.001). The Mele Bay (Vanuatu) population of *Nautilus vanuatuensis* exhibited the heaviest mean δ^13^C values (–13.18 ± 0.37‰), while the Great Barrier Reef population of *N. stenomphalus* showed the lightest (–17.20 ± 1.70‰). Although these represent the extremes across all sampled populations, they are not directly comparable at the species level due to their geographic endemism. Nevertheless, when comparing species overall, we detected a significant difference in δ^13^C values among taxa (F_4,76_ = 26.04, p < 0.001).

Pearson correlation analysis revealed a statistically significant but biologically negligible correlation between δ^13^C and shell diameter in the GBR population (r^2^ = 0.0029, p = 0.79), and slightly stronger correlations in the Bohol Sea (r^2^ = 0.44, p = 0.001) and Mele Bay (r^2^ = 0.47, p = 0.08). Although the Bohol Sea population shows statistical significance, the strength of these correlations suggests limited biological relevance. Notably, a loess curve revealed a slight dip in δ^13^C in intermediate-sized individuals followed by an increase in larger nautiloids.

### Values of δ^15^N

There was significant difference among the different nautiloid populations (F_6_,_93_ = 68.74, p<0.001). The values were highest in *Nautilus* found in the pooled Papua New Guinea (Kavieng and Ndrova) populations (15.88 ± 0.5 ‰) and lowest in the Fiji *Nautilus* population (10.71± 0.42‰). In terms of species of *Nautilus*, there was significant difference among the different species (F_4_,_76_ = 24.34, p<0.001). The values were highest in *Nautilus samoaensis* (15.6 ± 0.32‰) and lowest in *Nautilus vitiensis* (10.71± 0.42‰).

We found a significant positive correlation between δ^15^N and shell diameter in the GBR (r2=0.05, p = 0.24) Bohol Sea (r2=0.23, p = 0.035) and Mele Bay (r2=0.4, p = 0.12), as illustrated in Figure 5. Our data indicates that larger adult nautiluses tend to have higher enrichment of δ^15^N.

### Isotopic niches of nautiloids

The isotopic niche represents a quantitative approximation of an organism’s trophic ecology, defined by the distribution of their Carbon and Nitrogen isotopic ratios. This multidimensional space reflects the range of assimilated resources. Isotopic niches differ significantly among regions, with considerable overlap between some sites (e.g., Beqa Harbor and Pago Pago), while other locations, such as the Bismarck Sea and Bohol Sea, show clear separation. Standard ellipse analysis [26] revealed that the narrowest niche was observed in American Samoa (TA = 0.24, SEAc = 0.3), based on both the convex hull area (TA) and the small-sample corrected standard ellipse area (SEAc), whereas the largest niche was found in the Great Barrier Reef (TA = 10.29, SEAc = 3.1). However, the results for American Samoa were based on only five specimens, which may have led to an underestimation of the niche size.

When observed in a species perspective (Figure 3), isotopic niches varied across *Nautilus* species and *A. scrobiculatus*, with notable differentiation in niche width and centroid position. *N. pompilius* populations, in regions where they are present, exhibited variability based on geographic origin (e.g., Philippines vs. Papua New Guinea), while *N. stenomphalus* and *N. vanuatuensis* have more constrained niches.

**Figure 1.**
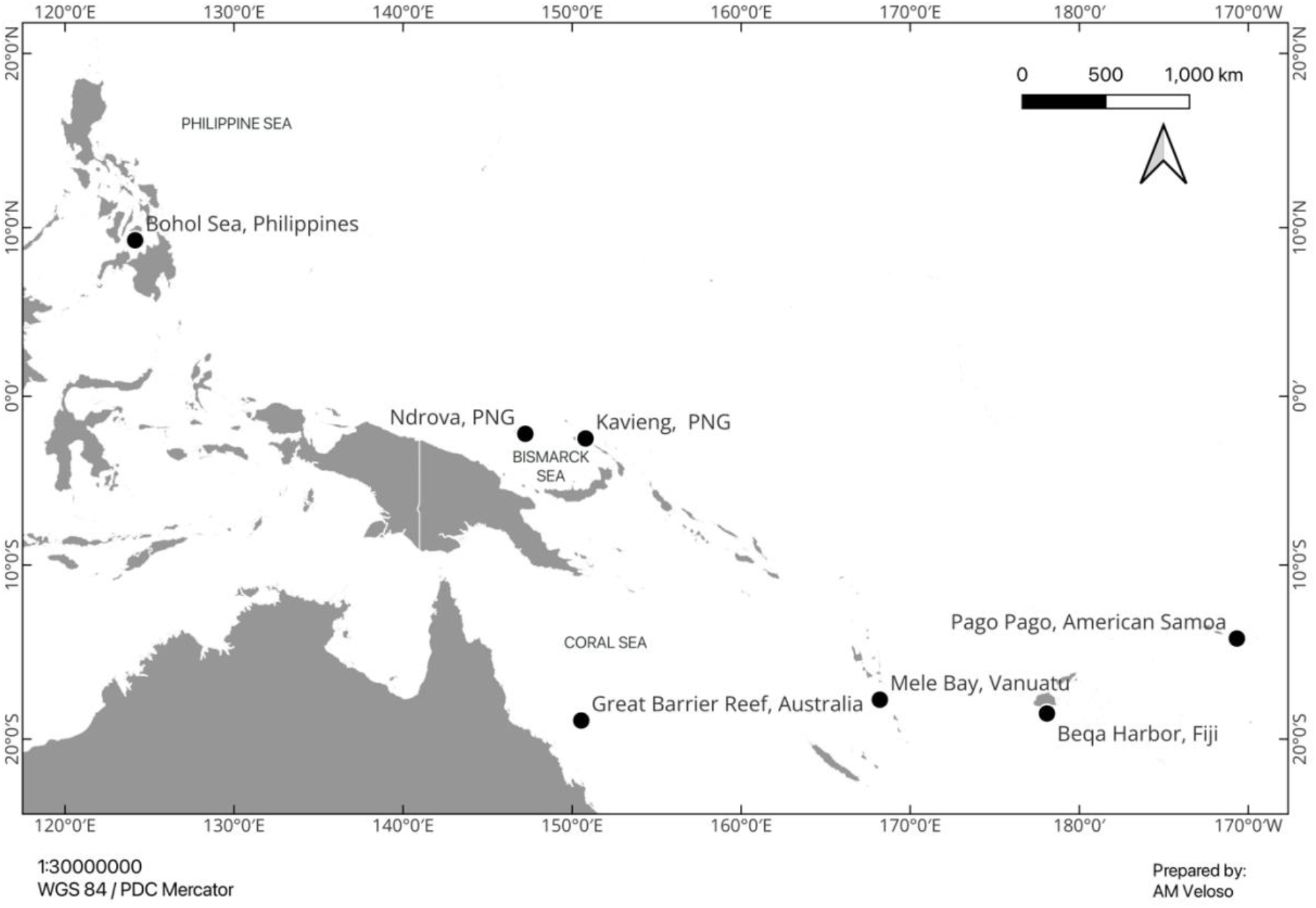
Nautiloid populations across multiple sites in the Indo-Pacific region, including the Philippines, Vanuatu, the Great Barrier Reef of Australia, Fiji, and American Samoa during the summer months of 2011 and 2012. Additionally, we collected samples from Kavieng and Ndrova in Papua New Guinea during May 2022. Available in QGIS.

**Figure 2.**
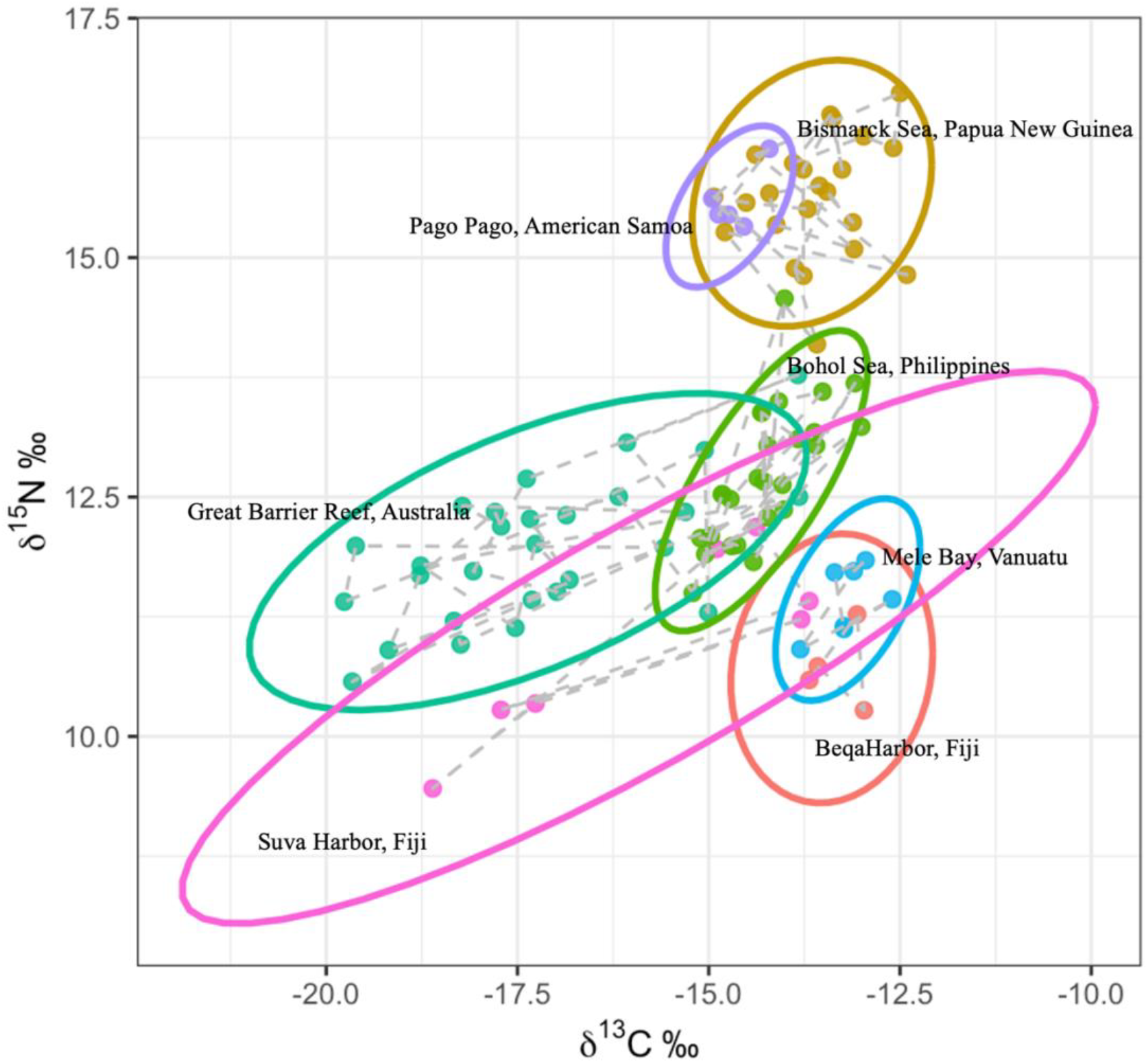
Stable isotope ellipses of δ^15^N and δ^13^C values for nautiloid populations around the Pacific Ocean. The standard ellipse represents the core isotopic niche of each nautiloid population; the convex hulls contain all data points from each nautiloid population.

**Figure 3.**
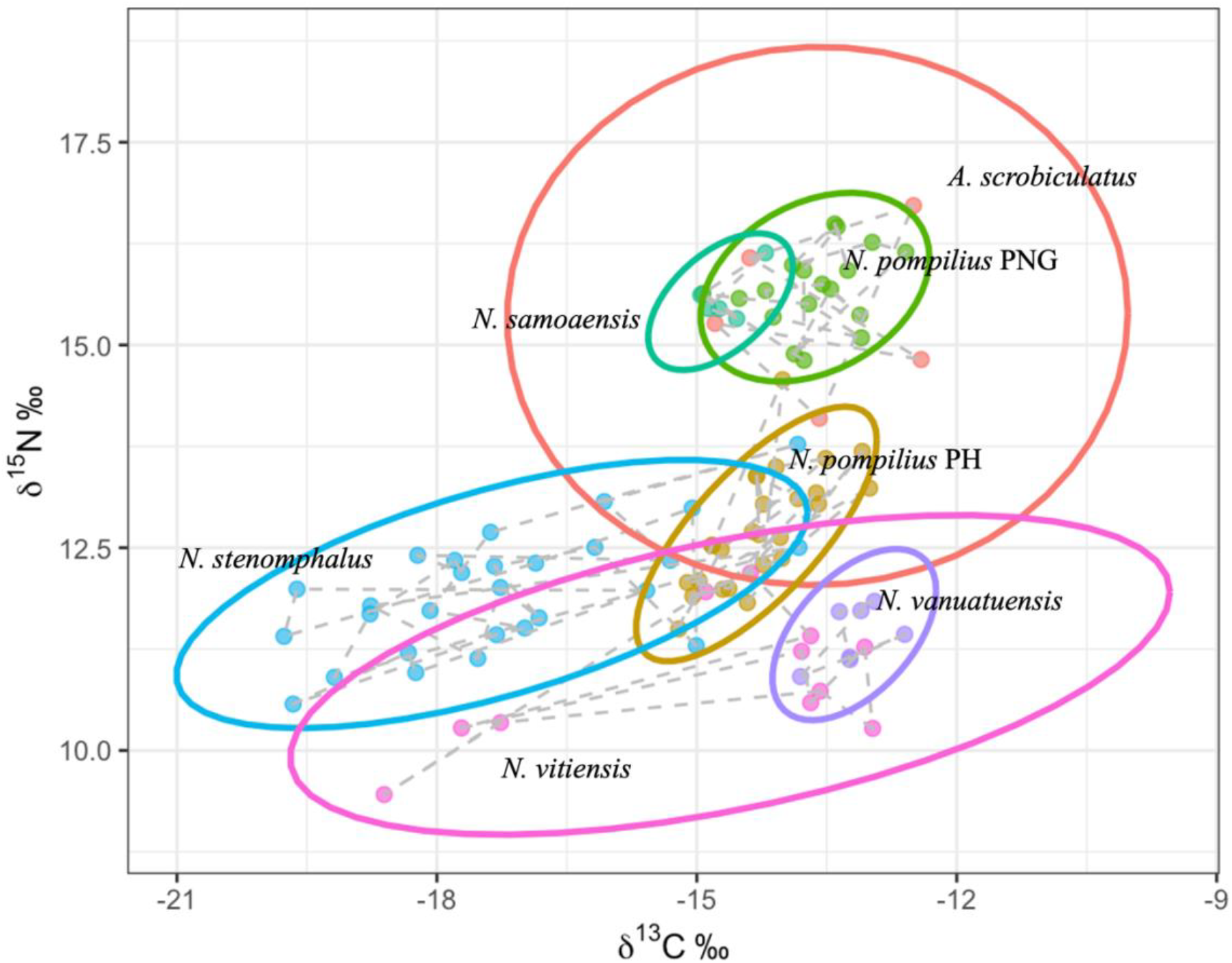
Isotopic niches of the nautiloid species presented in this study. Solid lines represent the ellipse area. The dotted lines represent the convex hull area.

**Figure 4.**
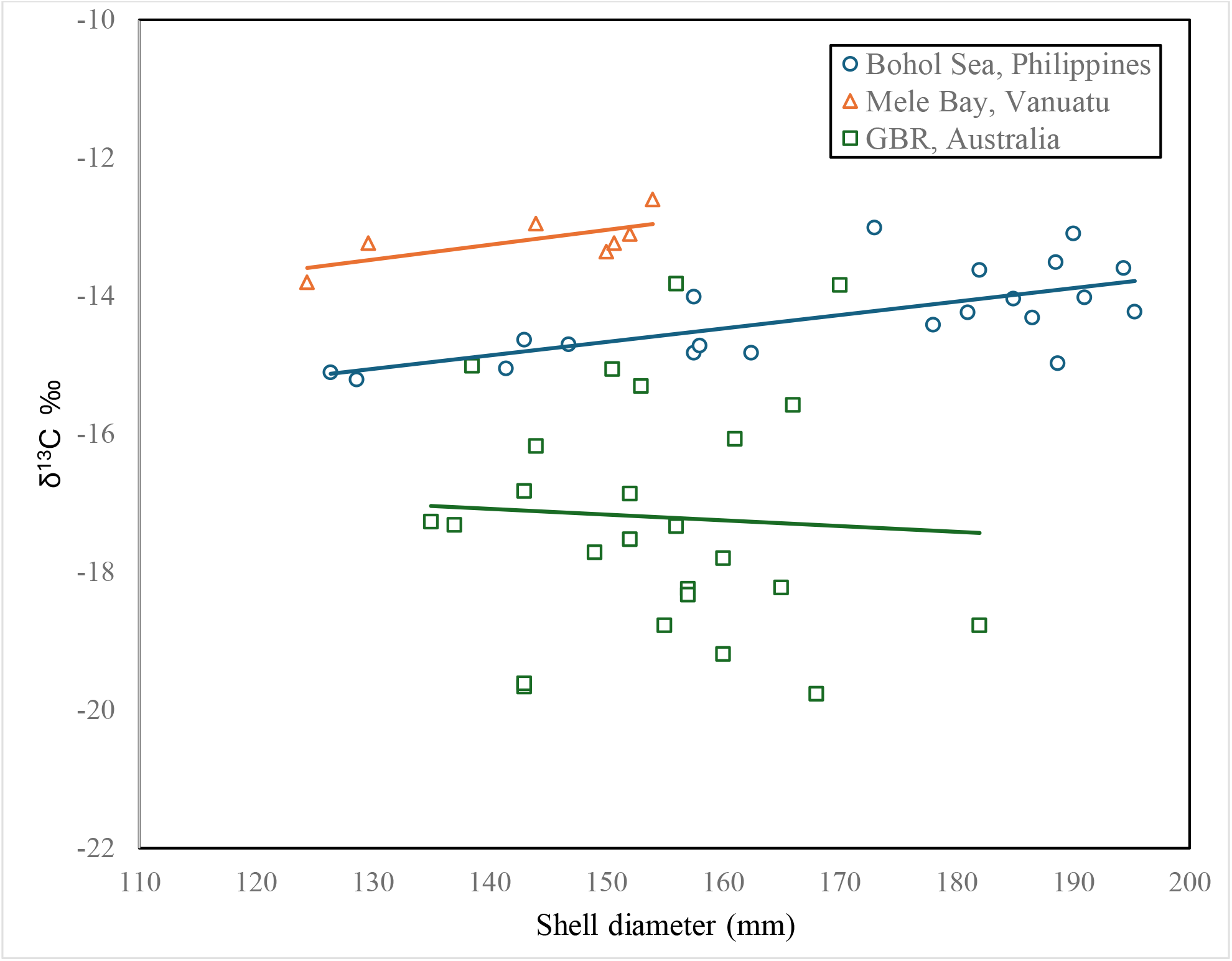
Relationship between shell diameter and δ^13^C for *Nautilus* populations from the Bohol Sea (Philippines), Mele Bay (Vanuatu), and the Great Barrier Reef (Australia). Each symbol represents an individual specimen, with trend lines indicating population-specific patterns. The Vanuatu population exhibits the highest δ^13^C values, while the GBR population has the lowest, suggesting regional differences in carbon source utilization or habitat preference.

**Figure 5.**
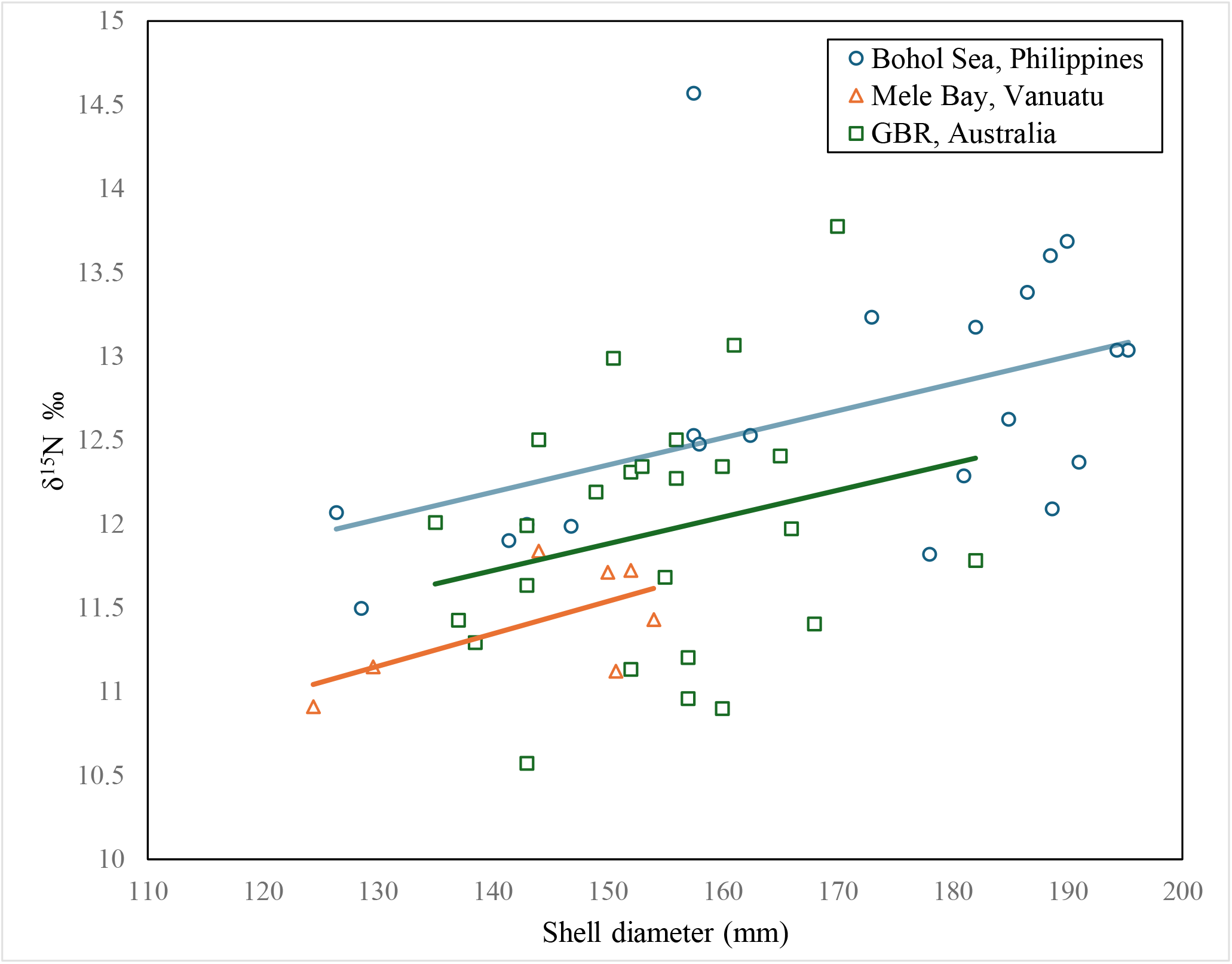
Relationship between shell diameter and δ^15^N for *Nautilus* populations from the Bohol Sea (Philippines), Mele Bay (Vanuatu), and the Great Barrier Reef (Australia). Each symbol represents an individual specimen, with trend lines indicating the direction and strength of the relationship. The Bohol Sea population shows relatively high δ^15^N values and the steepest slope, while the Vanuatu population has the lowest δ^15^N values and the shallowest slope, suggesting regional differences in trophic position or nitrogen baselines.

**Table 1.**
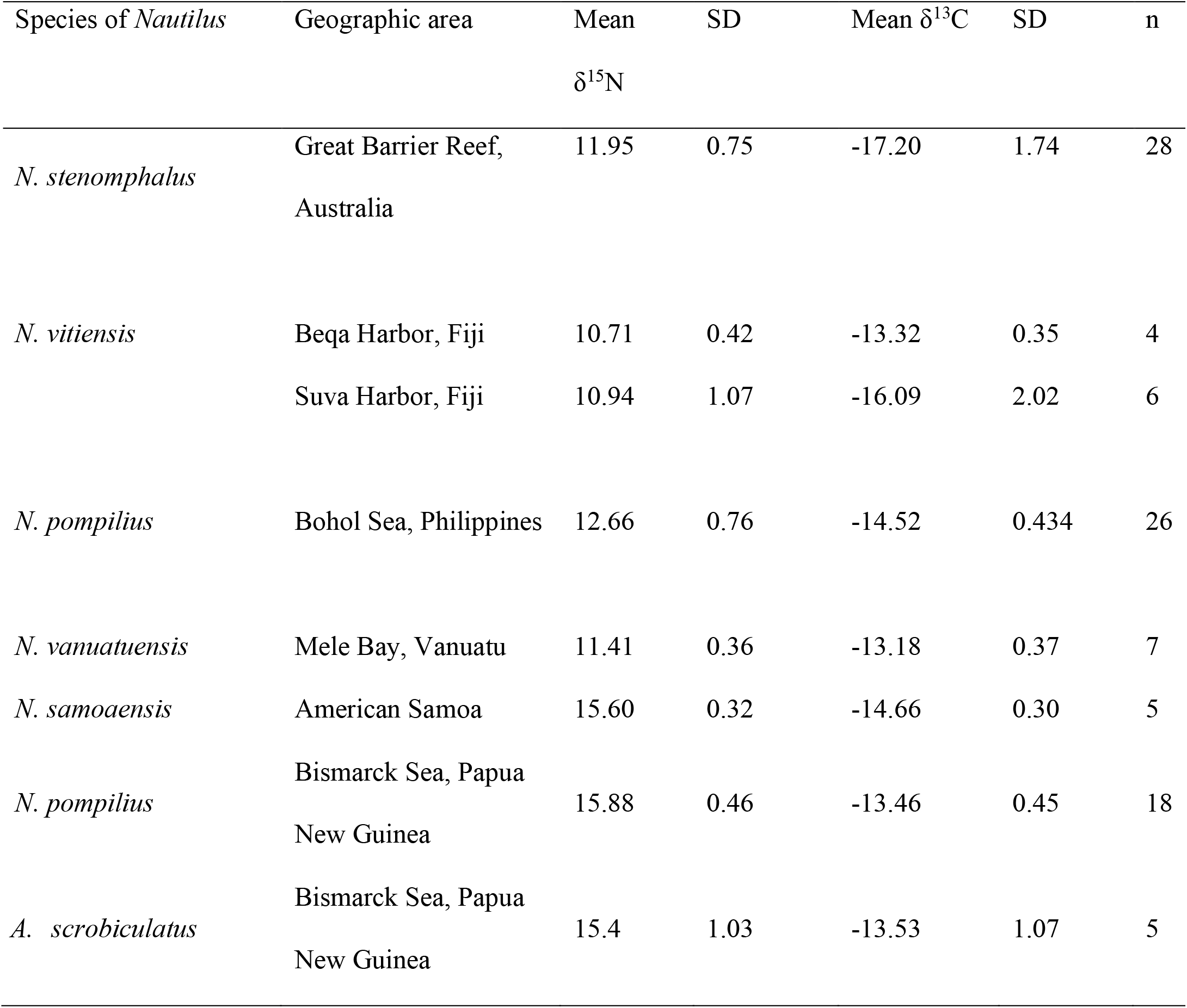
Summary of Carbon and Nitrogen isotope values categorized in species and locality.

Individual δ^13^C and δ^15^N values and associated shell measurements for each population are provided in Supporting Information (S1 Dataset).

## Discussion

Isotopic analyses reveal distinct ecological patterns among nautiloid populations across the Pacific, highlighting both unique and overlapping niche adaptations shaped by local biological and oceanographic conditions. Although niche spaces overlap across the three major clades [14]: the South Pacific clade (*N. samoaensis, N. vitiensis, N. vanuatuensis*), the Coral Sea clade (GBR and Papua New Guinea), and the Indo-Pacific clade (*N. pompilius* from the Philippines). Our results support the view that geographic setting, more than phylogenetic identity, governs isotopic niche differentiation. Distinct regional environments such as the Bismarck Sea or the GBR impose localized trophic dynamics, while overlaps among some populations suggest shared resource use or similar baseline isotope values [28,29].

The Great Barrier Reef provides a striking example. Here, *N. stenomphalus* exhibits a broad isotopic niche, which may reflect a diverse food availability and reef-derived carbon inputs [30–35]. In contrast, *N. vanuatuensis* in Port Villa shows a narrow niche, consistent with resource-limited systems [36]. Elevated δ^15^N values in the Bismarck Sea and American Samoa further illustrate how regional nutrient cycling, including processes such as nitrogen fixation, can shape local isotopic baselines [37,38]. In marine systems, spatial variation in baseline δ^15^N can be substantial, meaning that differences in consumer isotope values may reflect both trophic position and underlying environmental variation. As such, the observed differences in δ^15^N among nautiloid populations should be interpreted as reflecting regional isotopic niche structure, rather than direct differences in trophic level alone. This highlights the importance of local oceanographic context in shaping isotopic signatures in deep-sea organisms.

Ontogenetic trends add another dimension. In several populations, δ^15^N increased with shell diameter, reflecting dietary shifts toward higher-trophic prey as individuals matured [22,39–46]. This aligns with patterns reported in other cephalopods, though contrasts with septal isotope records in nautiloids that capture yolk-derived nitrogen inputs early in life [47]. By focusing on organic tissues from live-caught animals, our study captures present ecological behavior and highlights how foraging strategies shift across life stages, likely in response to prey availability, predation risk, and habitat use [45,48,49].

A key limitation of this study is the absence of region-specific baseline isotope measurements, which constrains our ability to fully disentangle variation in trophic position from underlying spatial differences in δ^15^N. In marine systems, baseline isotope values can vary substantially due to processes such as nitrogen fixation and regional nutrient cycling, meaning that differences in consumer δ^15^N may reflect both ecological and environmental drivers. As such, the patterns observed here are best interpreted as reflecting regional isotopic niche structure rather than direct differences in trophic level alone.

Taken together, our findings show that geographic setting appears to be a strong driver of isotopic niche differentiation, likely through a combination of environmental baseline variation and ecological processes influencing resource use.. Isolated populations exhibit unique isotopic signatures, some broad and flexible, others narrow and constrained. This geographic structuring may help explain how nautiloids persist across fragmented reef slopes, but also points to vulnerabilities where resource limitation narrows niche breadth. More broadly, our results highlight the ecological flexibility of this ancient lineage and suggest that spatial environmental variation, rather than phylogeny alone, plays a decisive role in shaping their trophic strategies and evolutionary trajectories.

## Acknowledgements and funding sources

This research was funded in part by Save the Nautilus, the National Science Foundation under Cooperative Agreement No. DBI-0939454 (BEACON: Evolution in Action), the Tiffany & Co. Foundation grant No. 11661, National Oceanic and Atmospheric Association grant No. NA12NMF-4690220, and by the US Fish and Wildlife Service (FWS) grant No. 10170-85-001.

## Author contributions

J.L.V. conceived the study, conducted data analyses, prepared all figures, and wrote the initial draft of the manuscript. P.D.W. contributed to conceptual development and writing. G.S.B. and F.D. assisted with fieldwork and provided minor text revisions. All authors reviewed and approved the final manuscript.

## Competing interests

The authors declare that they have no financial or non-financial competing interests related to this work.

## Notes

### Competing Interest Statement

The authors have declared no competing interest.

